# Notos - a Galaxy tool to analyze CpN observed expected ratios for inferring DNA methylation types

**DOI:** 10.1101/180463

**Authors:** Ingo Bulla, Benoît Aliaga, Virginia Lacal, Jan Bulla, Christoph Grunau, Cristian Chaparro

## Abstract

**Background:** DNA methylation patterns store epigenetic information in the vast majority of eukaryotic species. The relatively high costs and technical challenges associated with the detection of DNA methylation however have created a bias in the number of methylation studies towards model organisms. Consequently, it remains challenging to infer kingdom-wide general rules about the functions and evolutionary conservation of DNA methylation. Methylated cytosine is often found in specific CpN dinucleotides, and the frequency distributions of, for instance, CpG observed/expected (CpG o/e) ratios have been used to infer DNA methylation types based on higher mutability of methylated CpG.

**Results:** Predominantly model-based approaches essentially founded on mixtures of Gaussian distributions are currently used to investigate questions related to the number and position of modes of CpG o/e ratios. These approaches require the selection of an appropriate criterion for determining the best model and will fail if empirical distributions are complex or even merely moderately skewed. We use a kernel density estimation (KDE) based technique for robust and precise characterization of complex CpN o/e distributions without *a priori* assumptions about the underlying distributions.

**Conclusions:** We show that KDE delivers robust descriptions of CpN o/e distributions. For straightforward processing, we have developed a Galaxy tool, called Notos and available at the ToolShed, that calculates these ratios of input FASTA files and fits a density to their empirical distribution. Based on the estimated density the number and shape of modes of the distribution is determined, providing a rational for the prediction of the number and the types of different methylation classes. Notos is written in R and Perl.

## Background

### DNA methylation is an important bearer of epigenetic information

In eukaryotes, methylation occurs in the 5’ position of the pyrimidine ring of cytosine, leading to 5-methyl-cytosine (5mC), which can subsequently be converted into hydroxy-5-methyl-cytosine [6]. The presence of 5mC can have an impact on gene expression [48], alternative splicing [28] and other biological processes. Compared to other bearers of epigenetic information, such as posttranslational histone modifications and non-coding RNA, 5mC appears to be relatively stable and epimutation rates at this base rarely exceed 10^*-*^4 per generation [46]. The modification is also chemically very stable and survives common conservation methods for biological material. DNA methylation is therefore very often the target of choice when it comes to studying the impact of epigenetic information on the phenotype and the heritability of epiallels.

### DNA methylation and CpN o/e ratios

Several techniques are available to study 5mC distribution. Nevertheless, the relatively high costs of DNA methylation analyses have led to a bias in the results towards model organisms and towards the biomedical field. For the moment, it is not feasible to obtain comprehensive DNA methylation results for a large range of phylogenetic branches. This (i) is an obstacle to the introduction of epigenetics in fields in which historically the domain is not entirely accepted (e.g. ecology and evolution), and (ii) more importantly might lead to misinterpretation of results obtained in phylogenetically dissimilar (non-model) organisms. In many species, 5mC occurs either predominantly or exclusively in CpG pairs. This and the tendency of 5mC to deaminate spontaneously into thymine leads in methylated genomes to an under-representation of CpG over evolutionary time scales [26]. In molds, methylation can also be concentrated in CpA pairs and CpA o/e was used as an indicator of a process called repeat-induced-point-mutations (RIP) in which 5mC serves as mutagen, converting rapidly 5mC into thymine. Consequently, the ratio of observed to expected CpG pairs (CpG o/e) (and CpA o/e in fungi) was used to estimate the level of DNA methylation early on: in the methylated compartments of the genome, 5mCpN will tend to be mutated into TpN and the CpN o/e ratio will decrease. In contrast, in unmethylated genomes, the ratio will be close to 1. In principle, it is therefore conceivable to infer methylation in DNA on the basis of CpN o/e, and to do this for any species for which genome and/or transcriptome sequence data are available [54]. This could then provide a starting point for more detailed biochemical DNA methylation analyses. For many species, the available mRNA data outnumber largely the available genome sequences.

### Robust description of CpN o/e ratios is challenging

In the following study we will focus on mRNA even though the method we will describe can be used on any type of DNA/RNA sequence. For the sake of clarity, in this manuscript, we will also use primarily methylation in the CpG context, although our approach can be applied to any (multiple)nucleotide frequency distribution. Simple Gaussian distributions can be used in some cases to describe CpG o/e distributions. But in many species, methylation distribution is heterogeneous, leading to complex mixtures in CpG o/e distributions over all genes, and the Gaussian mixture approach will fail. Many invertebrates, for instance, possess a mosaic type of methylation with large highly methylated regions intermingled with regions without methylation [42]. To our knowledge, no method exists that allows for a straightforward data processing of CpG o/e for non-specialists that is usable for all types of CpG o/e data. Here, we describe such a tool that we called Notos. We tested Notos on all data available in dbEST [8] since this database is one of the most widely used and covers a wide range of species. Notos integrates into Galaxy but is also available as suite of stand-alone scripts, it requires little computational resources, and the analysis is done within minutes. It is thus suitable for the routine first-pass prediction of DNA methylation in many biological settings.

## Methods

Notos is a kernel density estimation (KDE) based tool. Its implementation is computationally efficient and allows for processing even large data sets on an ordinary personal computer. The analysis carried out by the Notos suite is composed of two steps and corresponds to two separate programs (see Figure 1 for the work flow): First, the preparatory procedure CpGoe.pl calculates the CpG o/e ratios of the sequences provided by a FASTA file. Any CpN o/e can be calculated if supplied as parameter. Secondly, the core procedure KDEanalysis.r, which consists of an R script [45] carrying out two principal parts: data preparation and analysis of the distribution of the CpG o/e ratios using KDE. It is also possible to skip the preparatory procedure and directly provide KDEanalysis.r with CpG o/e ratios - or other data of comparable structure. We describe the two steps in the following.

**Figure 1.**
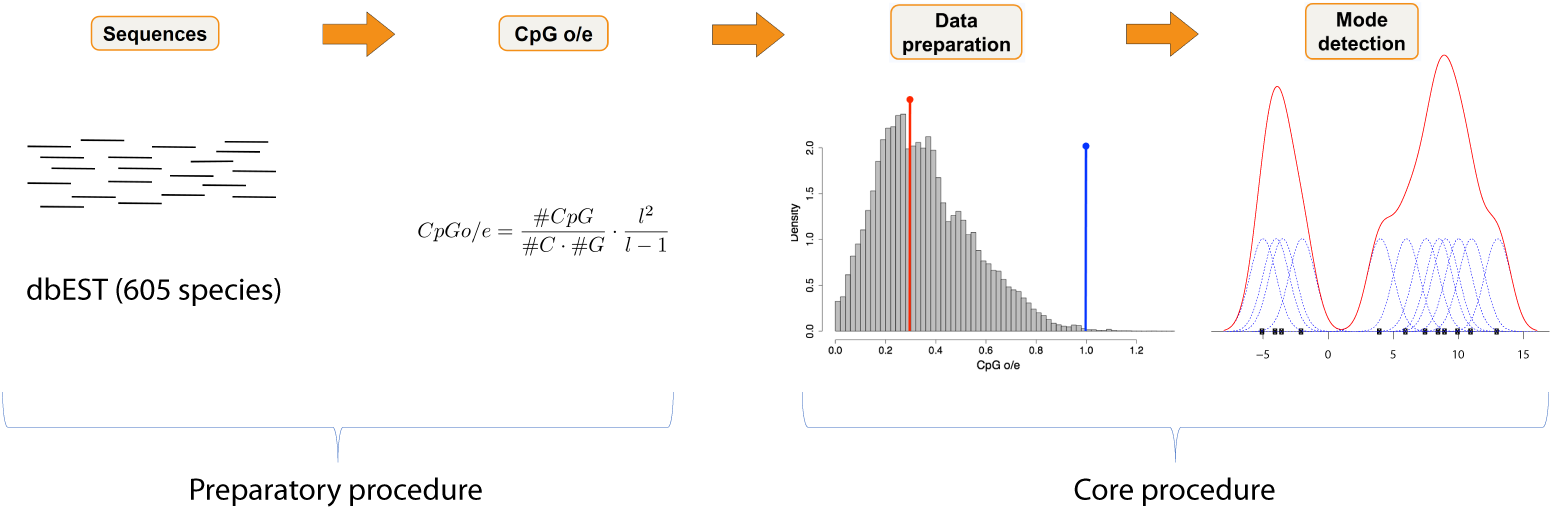
Workflow. Steps: 1. CpGo/e ratios are calculated for the sequences to be analyzed (in our case dbEST) using CpGoe.pl. 2. Removal of outliers (first step of KDEanalysis.r). 3. Mode detection (second step of KDEanalysis.r)

### Preparatory Procedure: Data Input

The data necessary as input for the core procedures of Notos are CpG o/e ratios in form of a vector. These ratios correspond, in principle, to the number of CpGs observed in a sequence divided by the number of CpGs one would expect to observe in a randomly generated sequence with the same number of cytosine and guanine nucleotides.

#### Literature formulas

Several formulae for calculating this ratio have been established in the past years, all deriving some form of normalized CpG content. The presumably most popular versions (see, e.g., [32] and [20], respectively) are

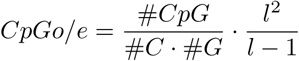

and

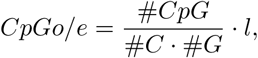

where *l* is the length of the sequence, and #*C*, #*G*, and #*CpG* denote the number of C’s, G’s, and CpG’s, respectively observed in the sequence. Alternative formulations were, among other, given by [56] who proposed

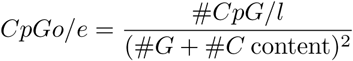

and by [36] with

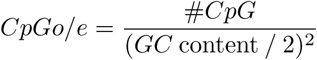

In their version, the #*G* + #*C* content is defined as the total number of C’s and G’s divided by the total number of nucleotides, and *GC* content is defined as the total number of C’s and G’s.

Notos The script CpGoe.pl allows the calculation of CpG o/e ratios from a multi-FASTA sequence and uses the formulation of [32] (i.e. the first formula above) by default, the others are optional. Moreover, sequences having less than 200 unambiguous nucleotides are eliminated from the calculation in the default setting, since our test runs indicated that too short sequences led to large amount of zeros or other extreme values.

## Core Procedure: Data Cleaning and Analysis via KDE

The core procedure KDEanalysis.r carries out two steps: first, data preparation, which is mainly necessary to remove data artifacts, and secondly mode detection via KDE. Both steps return the user results in form of CSV files and figures. In addition, they allow overriding the default settings, if this is required by the user. Note, however, that such changes should be carried out with care, since all settings have been calibrated through intensive testing procedures on several hundred species from the dbEST database. In the following paragraphs, we describe these two steps in detail.

### Data preparation

The first step, data preparation, starts by removing all values equal to zero from the input data since these observations correspond to artifacts resulting from too short sequences or sequences that do not present any CpG dinucleotide. Then, extreme and outlying observations are removed, i.e. all values outside the interval [*Q*25 *- kIQR, Q*75 + *kIQR*], where *Q*25, *Q*75, and *IQR* denote the 25% quantile, the 75% quantile, and the interquartile range, respectively. In order not to exclude too many observations, the threshold parameter *k >* 1 takes the smallest integer value ensuring that not more than 1% of the data are removed, whereby *k* cannot exceed the value five. We determined the value of 1% through testing on a large number of species, and found it to be a good compromise between the need to exclude as many outliers as possible and not changing the distributional properties of a sample in a substantial way.

The output of this step consists of a table with various summary statistics in CSV format, and a figure displaying the data before and after this step. Figure 2 corresponds to the output resulting from an arbitrarily selected species, the locust *Locusta migratoria*. The content of the resulting table is described in detail in the documentation of Notos, which can be found in the readme file or the help section of the galaxy interface. Additional files 1 and 2 contain results from this step for 603 species from dbEST.

**Figure 2.**
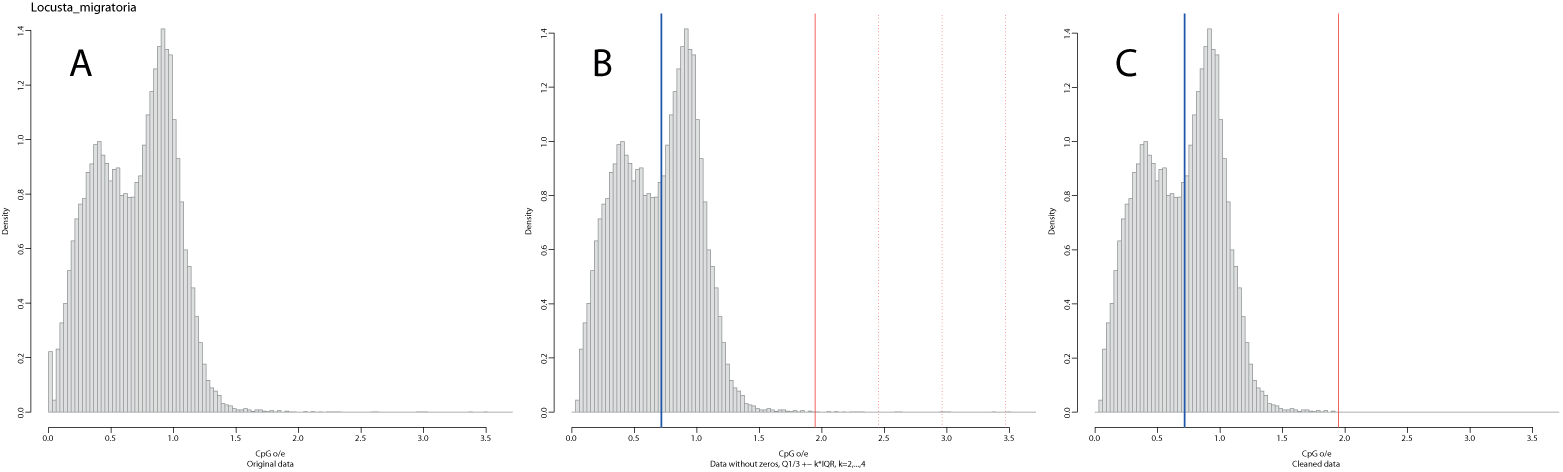
Step 1: data cleaning of a sample of CpG o/e ratios from the locust *Locusta migratoria*. The left panel (A) shows the original data. The middle panel (B) displays the data after removal of all values equal to zero. The blue vertical line corresponds to the sample median. Red vertical lines indicate the possible thresholds for excluding outliers and extreme observations. The selected threshold (*k* = 2) is solid, alternative thresholds are dotted. The right panel (C) shows the cleaned data with the sample median and the selected threshold.

### Mode detection

#### KDE

In the second step, we determine the number of modes by means of a KDE based procedure. The underlying statistical theory is well-established, and therefore described only briefly. In principle, it is assumed that the independent and identically distributed observations *x*_1_*, x*_*n*_*,…, x*_*n*_ constitute a sample with unknown density *f*.

Then, the kernel density estimator 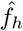 of *f* is given by

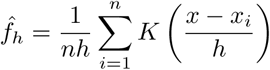

where *K*(.) is the so-called kernel function. The kernel function is non-negative, has a mean value equal to zero, and the area under the function equals one, i.e., *K*(.) satisfies the condition 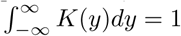. Several families of kernel functions are available, and we considered the most common ones (Gaussian and Epanechnikov) for the implementation of our algorithms. Finally, we selected the probably most common Gaussian kernel function with *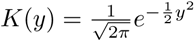* due to the satisfactory results obtained in practice. In order to determine the value for the smoothing parameter, which is commonly termed bandwidth as well, we investigated different possible approaches, such as cross-validation, Silverman’s rule [40], and Scott’s variation of Silverman’s rule [37]. Extensive testing on a large variety of species from different data sources suggested that the well-established bandwidth proposed by Scott provides the best results in terms of interpretability. Our bandwidth choice was motivated by the good interpretability of results obtained from testing on various data sources. In particular, it showed a satisfactory stability for species with either a very high or a very low number of observations.

#### Number of modes

Subsequently, the number of modes is then determined by counting the number of local maxima of the estimated density, and a probability mass is assigned to each mode. The calculation of this probability mass is straightforward by integrating the density over the interval determined by the next-nearest local minima to the left and right, respectively, of the mode. If no local minimum is present to the left (right), the integration limits are set to minus (plus) infinity. The resulting probability masses for all modes sum up to one, and provide a single value which serves, roughly speaking, for determining the importance of a mode. Last, the obtained results are post-processed by a) merging modes that are closer than 0.2 (default value) to each other and b) removing modes that accumulate less than 1% (default value) of the probability mass of the estimated density. Multiple peaks suggest multiple sequence populations with different methylation types. The rational behind step a) is that very close modes reflect very similar types of methylation and hence probably have no biological significance. The value of 0.2 as minimum CpG o/e distance was empirically determined based on organisms with known mosaic-type methylation and double CpG o/e modes. We believe that relying entirely on confidence intervals is not a valid option for species with very high numbers of observations and as a consequence narrow confidence intervals. The choice of the probability mass threshold of 1% for step b) resulted again from extensive testing on a large number of species. A mode with 1% or less of probability mass lying outside of the core part of the density would most likely result from contamination. An optional feature of the KDE analysis is the estimation of confidence intervals for the position of the modes as well as confidence estimates for the number of modes. This is implemented through case resampling (non-parametric) bootstrap with 1,500 repetitions. Since this part is slightly computationally demanding, the bootstrap is optional and is accelerated by parallel execution via the doParallel package.

#### Output

Similarly to the first step, the script KDEanalysis.r returns a figure to the user. Figure 3 shows this graphical output for the four species *Locusta migratoria*, *Alligator mississippiensis*, *Antheraea mylitta*, and *Citrus clementina*. The top panel A with *L. migratoria* shows two clearly distinct modes (blue vertical lines), their corresponding confidence intervals (shaded blue), and the fitted density (red). Moreover, a thin black vertical line indicates a local minimum, which serves for separating the probability masses attributed to each mode. In the case of *A. mississippiensis* (panel B), only one mode is present. Note that the confidence interval is strongly skewed, which results from the skewed empirical distribution used for the parametric bootstrap. For *A. mylitta*, one can observe that one of the two modes is assigned less than ten percent of probability mass, indicated by the dashed vertical line for the left mode in panel C. Last, *C. clementina* (panel D) possesses two modes relatively close to each other, i.e., the distance lies below the above mentioned threshold of 0.2. For this reason, the two modes may be interpreted as being too close for indicating biologically relevant differences in methylation types, which is underlined by their orange color. For results concerning other species from dbEST, see additional file 3.

**Figure 3.**
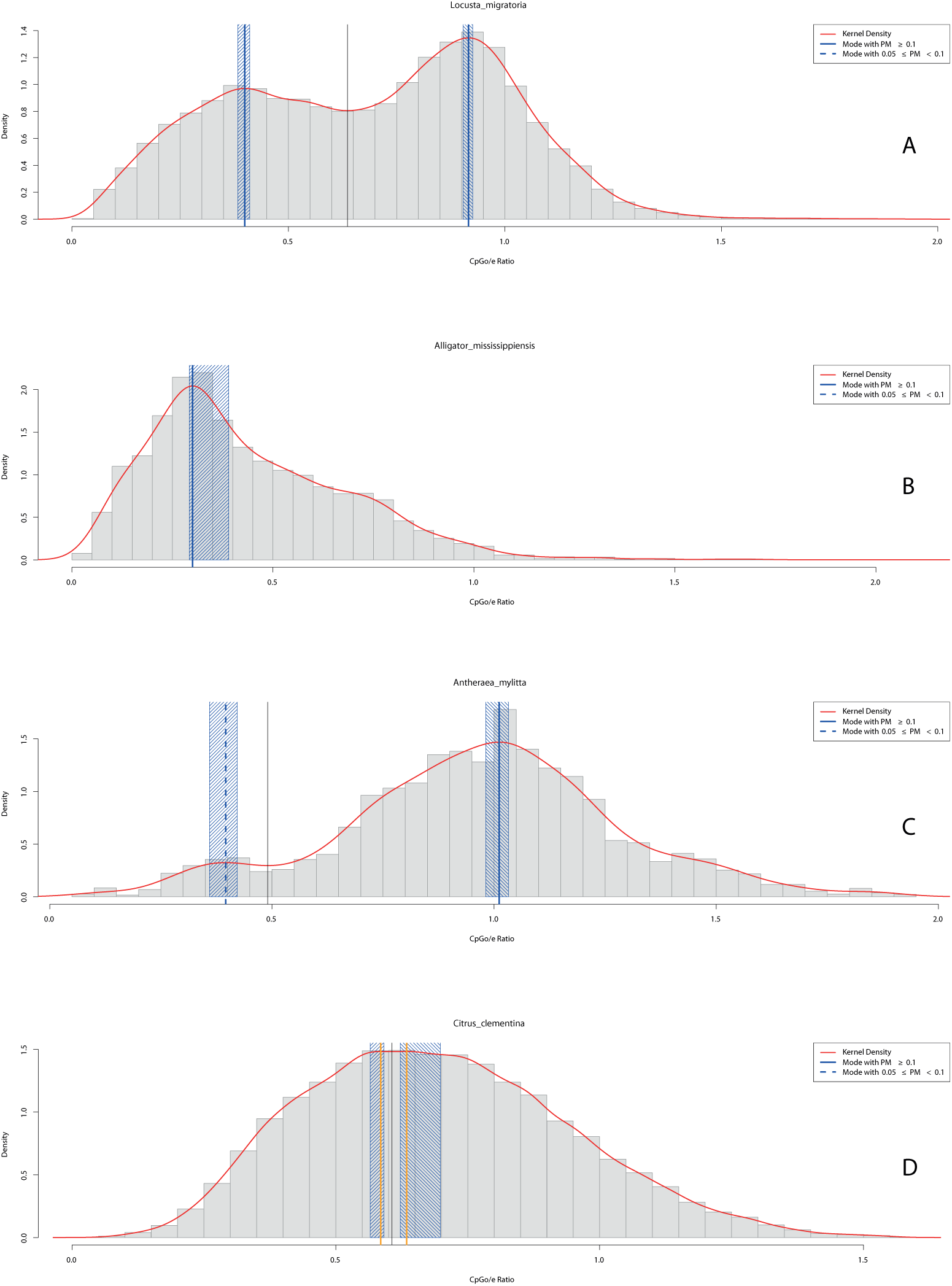
Step 2: kernel density estimation for samples of CpG o/e ratios from four species. The red line corresponds to the density estimated via KDE. Full vertical blue lines indicate modes with PM ≥ 0.1. Shaded blue areas around the modes correspond to bootstrap confidence intervals with a default level of 95%. From top to bottom, the panels show results for *Locusta migratoria* (A), *Alligator mississippiensis* (B), *Antheraea mylitta* (C), and *Citrus clementina* (D).

Moreover, the user obtains one table with various statistics related to the modes and their probability masses (see additional file 4 for the results for 605 species from dbEST). Optionally, a second table linked to the results obtained from the bootstrap procedure is generated (cf. additional file 5). The content of these two tables is also described in detail in the readme section of the Galaxy interface. The output from the bootstrap procedure deserves two additional remarks. Firstly, from a practical perspective, the number of modes identified in the bootstrap samples allows insight into the stability (and potential instability) of the number of identified modes. For example, at least one of the modes detected in the original sample should be considered weakly developed if a high proportion of bootstrap samples possesses a lower number of modes than the original sample. Alternatively, a frequently occurring higher number of modes in the bootstrap samples than in the original sample indicates that additional modes could develop with an increasing sample size - however, an increasing sample size may also have the opposite effect. Secondly, from a technical perspective, it may be non-trivial to assign modes identified in a bootstrap sample to the corresponding modes from the original sample, e.g., if several weakly developed modes are present in the original sample. In order to obtain reliable confidence intervals, two safeguards are implemented. On the one hand, bootstrap samples having a different number of modes than the original sample are excluded. On the other hand, samples with modes subject to strong changes (default value: 20%) in the probability mass compared to the original sample are excluded as well.

### Implementation

A Galaxy package has been created that allows the automated installation of the Notos suite in a Galaxy server. The suite installs an interface for CpGoe.pl which provides the calculation of the CpG o/e ratio as well as an interface for KDEanalysis.r which calculates the distribution of CpG o/e ratios using KDE. Empirical testing showed that at least about 500 sequences are necessary to obtain a reliable parametrization of the KDE for CpG o/e frequency distributions.

## Results

The test of Silverman [40] constitutes a classical, popular way to investigate multimodality. In the context of DNA methylation patterns, model-based approaches essentially founded on mixtures of Gaussian distributions have become a very popular approach to investigate questions related to the number of modes or underlying sub-populations [21, 18, 12, 5]. This popularity may result, inter alia, from the easy accessibility of statistical software allowing the treatment of mixture models, such as flexmix, mclust, or mixtools [4, 19, 27]. While the test of Silverman provides a rather simple criterion in form of a p-value rejecting (or not) the null hypothesis of a certain number of modes, model-based approaches require the selection of an appropriate criterion for determining the best model. The most prominent among established criteria are, e.g., the Akaike Information criterion (AIC) and its extensions, the Bayesian information criterion (BIC), and the Integrated Completed Likelihood (ICL) [see, e.g., 7, 10, and the references therein].

### Comparison

We investigated the performance of the Silverman test, the different criteria, and Notos on our data base with 603 species from dbEST. Table 1 shows the results from 17 arbitrarily chosen species, which display patterns that are representative of the full sample. The principal results are the following:

**Table 1.** This table shows the number of modes selected by different approaches and methods for 17 selected species: the test of Silverman (2nd column), model-based approaches, based on the criteria AIC, BIC, and ICL (3rd to 5th column) and Notos (last column). The maximum number of modes is limited to ten, all mixture models were estimated by the R-package mclust.

| Species | Silv. | AIC | BIC | ICL | Notos |
| --- | --- | --- | --- | --- | --- |
| *Acropora palmata* | 1 | 10 | 5 | 1 | 2 |
| *Actinidia chinensis* | 1 | 8 | 8 | 1 | 1 |
| *Aegilops speltoides* | 1 | 7 | 2 | 1 | 1 |
| *Aiptasia pallida* | 2 | 6 | 3 | 1 | 2 |
| *Alligator mississippiensis* | 1 | 7 | 4 | 1 | 1 |
| *Antheraea mylitta* | 4 | 6 | 3 | 1 | 1-2 |
| *Aspergillus oryzae* | 1 | 5 | 1 | 1 | 1 |
| *Bombus terrestris* | 2 | 5 | 3 | 1 | 2 |
| *Citrus clementina* | 1 | 8 | 4 | 1 | 1-2 |
| *Citrus limon* | 1 | 5 | 3 | 1 | 1 |
| *Danio rerio* | 1 | 8 | 8 | 1 | 1 |
| *Daphnia pulex* | 1 | 9 | 5 | 1 | 1 |
| *Drosophila melanogaster* | 2 | 8 | 3 | 1 | 1 |
| *Locusta migratoria* | 2 | 9 | 9 | 1 | 2 |
| *Nematostella vectensis* | 2 | 9 | 6 | 1 | 2 |
| *Pinctada maxima* | 1 | 10 | 4 | 1 | 2 |
| *Rattus norvegicus* | 1 | 10 | 8 | 1 | 1 |

i. The test of Silverman selects a low number of modes in most cases, with a few exceptions where the number of modes reaches high values. Overall, the number of detected modes is often difficult to explain or confirm by visual inspection of the sample, and the biological interpretation is (very) limited. Furthermore,
ii. The model selection criteria AIC and BIC generally produce non-interpretable results: both criteria allow for models with too many parameters, which regularly results in the selection of models with a far too high number of modes and no biological interpretability. This effect is illustrated in panel A of Figure 4 which shows the fitted density and the location of the component-specific means for *L. migratoria*, determined by the AIC solution. The discrepancy between the relatively clearly visible bimodal shape and the selected model with nine components is rather large. This high number of modes results from the very good fit to the empirical density for this sample containing a high number of observations. Panel B of Figure 4 illustrates the non-satisfactory performance of the BIC by means of *A. mississippiensis*. This species shows a single, clearly pronounced mode at approximately 0.3, and is strongly skewed to the right. This strong skewness leads to the additional identification of two components at about 0.6 and 1.0. Moreover, an additional component is identified at *∼*0.15 for compensating for another small deviation from normality.
iii. This drawback cannot be overcome by selecting the number of modes based on the ICL. This criterion almost always determines a single mode, which is sensible from a clustering perspective, but not desirable for mode identification, as panel C of Figure 4 shows.

**Figure 4.**
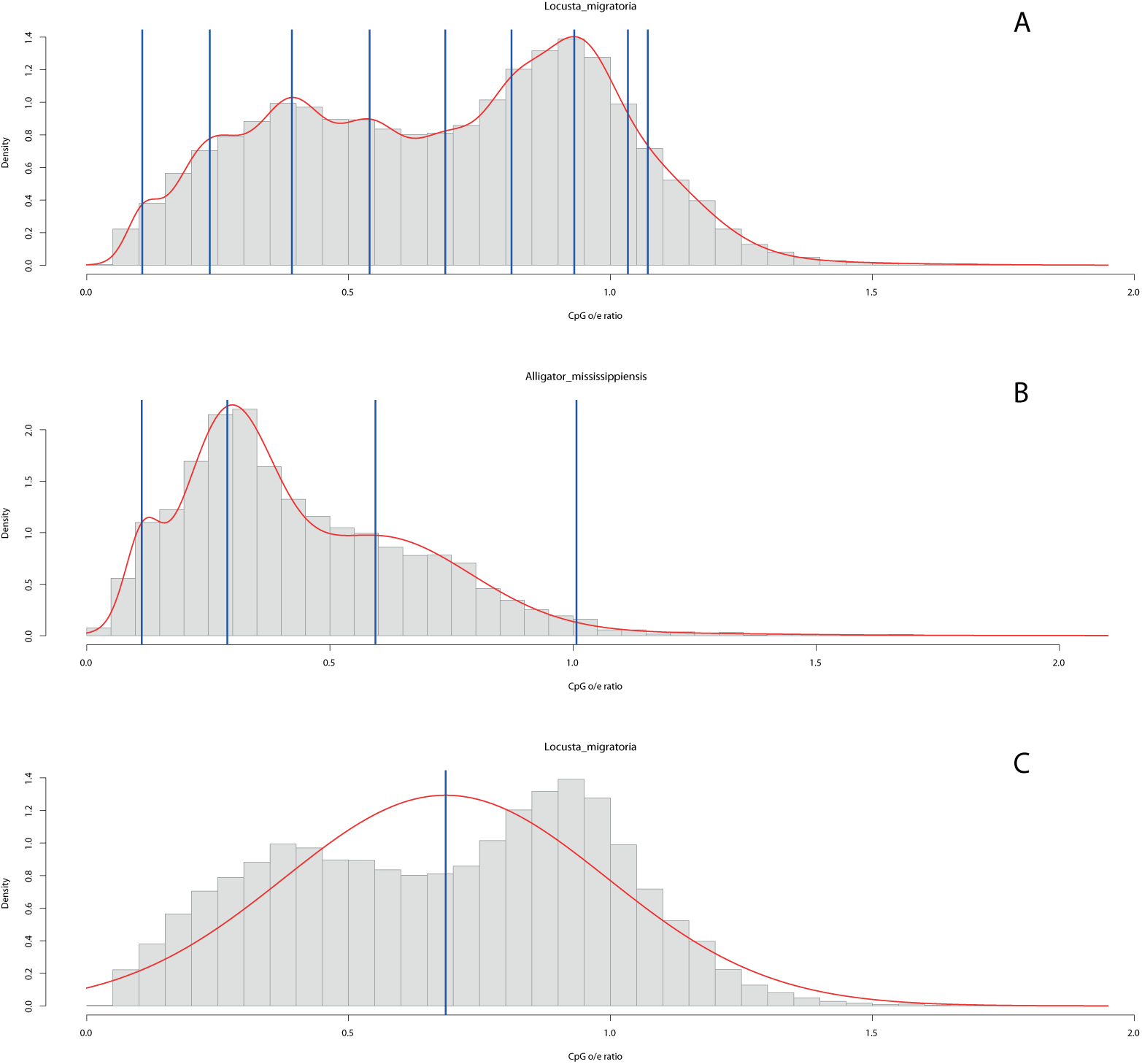
Examples for model-based clustering and model selection with Gaussian mixtures of CpG o/e ratios. The red line corresponds to the estimated density via KDE. Full vertical blue lines indicate the location of means belonging to each component of the mixture distribution (estimated by the R-package mclust). The top panel (A) shows the model selected by the AIC for *Locusta migratoria*, while the lowest panel (C) displays the corresponding ICL solution. The middle panel (B) displays the model selected by the BIC for *Alligator mississippiensis*.

### Interpretation

In conclusion, while conventional methods can perform well in many cases, they will also often fail to produce biologically interpretable results. For the 603 species from dbEST, the information criteria mentioned above as well as the test of Silverman fall short for approximately 60% of the data in this regard. In contrast, Notos performed well with all tested data sets. After having firmly established that Notos provides robust descriptions of mode locations and mode numbers, we attempted to establish a link between these parameters. As outlined above, a CpG o/e ratio around 1 is assumed to occur in non-methylated sequences and a ratio below 1 in methylated sequences. Consequently, if both situations are detected, both types of sequences co-exist in the studies sequence population. Based on comparison of Notos results with available literature data on DNA methylation, we tentatively assigned a threshold value of 0.85 to differentiate presumably methylated (*<*0.85) from presumably non-methylated (≥0.85) sequences. This is slightly higher than the 0.6, conventionally used e.g. for the detection of generally unmethylated CpG islands [25]. Further, more detailed analyses will be necessary to define parameters more precisely and to achieve a more detailed characterization of CpG o/e types and corresponding DNA methylation types.

### Case studies

To illustrate the use of Notos in two CpN contexts, we will present in the following results for the classical model species *Neurospora crassa*. *N. crassa* is a mold that belongs to the ascomycota. DNA methylation in this species is well described: only repetitive sequences such as relicts of transposons but not protein coding genes are methylated [39]. Methylation in these regions is associated with a genome defence system called repeat-induced point mutations (RIP) (reviewed in [38]). This system targets specifically CpA dinucleotides [11] where C is converted into T. CpA depletion is considered as a sign of RIP in other fungal species as well [13]. We therefore anticipated that CpG o/e and CpA o/e ratios in coding sequences would be around or above 1 (no methylation), while CpA o/e ratios, but not CpG o/e ratios, would be clearly below 1 in repeats indicating methylation in this context. We used the Neurospora crassa.ASM18292v1.31.dna sm.genome.fa genome assembly and the corresponding Neurospora crassa.ASM18292v1.31.gff3 annotation file from http://fungi.ensembl.org/Neurosporacrassa/Info/Index to extract 40,826 sequences for repeats and 10,432 sequences of spliced exons. A minimum length of 1 kb was used. As expected, a distribution with a single mode at a maximum at 0.9-1.1 was observed for CpG and CpA o/e ratios in spliced exons (panel A and B, respectively of Figure 5). In contrast, the mono-modal CpA o/e ratio distribution in repeats peaked at 0.47, while for CpG o/e the single mode was shifted towards 1.5 (panel C and D of Figure 5). The results of this straightforward and rapid analysis correspond therefore entirely to what is known about DNA methylation in *N. crassa*.

**Figure 5.**
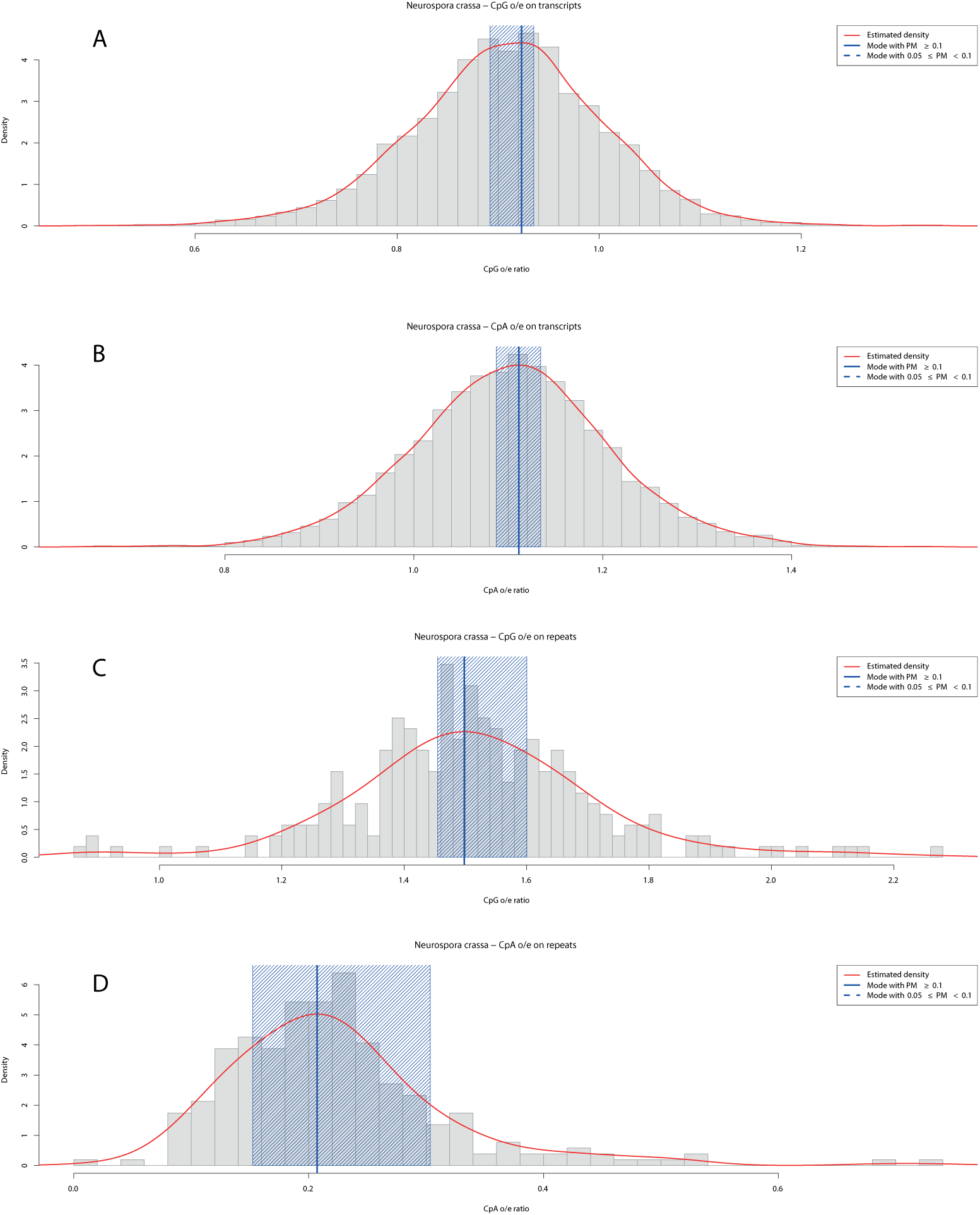
CpN o/e analyzed by Notos for *Neurospora crassa*. The red line corresponds to the estimated density via KDE. Full vertical blue lines indicate modes with PM ≥ 0.1. Shaded blue areas around the modes correspond to bootstrap confidence intervals with a default level of 95%. The panels show kernels of transcripts for CpG o/e (A) and CpA o/e (B), and for repeats (C and D), respectively. In this case CpG and CpA o/e ratios were calculated for spliced exons and repeat regions of the *N. crassa* genome. Both o/e frequency distributions are clearly unimodal, but for the CpA o/e in repeats there is a shift towards 0.5 which is concordant with DNA methylation only in this context (repeats and CpA) in this species.

## Discussion

DNA methylation is a conserved feature of many genomes. Since it remains neutral in its protein coding potential its use for adding additional epigenetic information to the DNA has been evolutionary stable. Nevertheless, the type of encoded information and consequently the type of DNA methylation can vary considerably, and many species have no or very little DNA methylation. It is thus of great practical value to be able to propose a well-founded hypotheses or at least educated guess about the type of methylation in a biological model before choosing an experimental strategy to study it in more detail. Notos was generated to produce such testable hypothesis.

### Technical alternatives

It could be argued that other wet-bench based methods deliver comparable results about the presence and the type of methylation. It is for instance straightforward to digest DNA with methylation sensitive restriction enzymes [31] and to separate the resulting fragments by electrophoration. A digestion smear would indicate absence of methylation. But this requires producing sufficient amounts of high-quality DNA, which is not always possible (e.g. protected or rare species, degraded DNA, samples that are difficult to obtain). Digestion is also difficult to quantify. Extensions of the digestion method are methylation sensitive amplified length polymorphism (MS-AFLP) [53], reduced-restriction bisulfite sequencing (RRBS) [33] or reference-free reduced representation bisulfite sequencing (epiGBS) [47]. These methods are very powerful and can be used with or without a reference genome (that is not necessarily available for non-model species). A caveat of RRBS is however that it was designed for the methylation type of vertebrates that typically possess methylation free CpG islands. It might not work well with other methylation types. Similarly to the simple digestion method, all these methods need physical access to high quality DNA and require already considerable investment (currently from several hundreds to thousands of euros). The same applies for more exhaustive and more expensive affinity based methods (such as MeDIP) [50] or whole genome bisulfite sequencing (WGBS) [29]. In many cases, a biochemical analysis of DNA methylation will hence be difficult and would require time and labor-intensive acquisition of DNA as well as investment in optimization of the analysis. Especially researchers with little biomolecular knowledge will hesitate to engage in investigations on DNA methylation even though they possess a perfect expertise about their species of interest and epigenetic insights would present advancements to them. These technical difficulties have led to a distortion in the available methylation information. A review of the available data in databases and in the literature showed that at least 300 methylomes are available for Human, mouse and the model plant *Arabidopsis thaliana* but only 63 for a total of 16 other species [3, 15, 30, 55, 23, 17, 9, 24, 22, 52, 51, 16, 49, 35, 1, 44, 43, 34, 14].

### Gaussian mixtures

When analyzing CpG o/e ratios related to DNA methylation, the model selection criteria AIC and BIC are regularly used for determining whether a model with two Gaussian components should be preferred to a simple normal distribution. This approach is at least questionable for two reasons. Firstly, model selection should be carried out taking a large number of possible models into account, and not just two (conveniently) selected alternatives. In our setting, it seems natural to consider models with more than two components as well, since the restriction to one or two components seems hard to justify from a biological perspective. This leads, however, to solutions that are (very) difficult to interpret. Secondly, models with two components may describe entirely different phenomena: on the one hand, the second component may result from a well-developed second mode. On the other hand, the second component may just result from minor deviations from normality, such as skewness or excess kurtosis. The latter behavior of both criteria results from the tendency to provide a good fit of the estimated density to the empirical data and put less emphasis on the clustering aspect, a fact investigated in more detail, e.g., by Baudry *et al.* [2].

### Other approaches investigated

Investigating confidence intervals and their properties (width, overlap) may provide additional insight, but requires a case-by case investigation which may then lead to subjective conclusions. We also tried to find a better balance between mode (or component) identification and non-normality by fitting mixtures of non-Gaussian distributions, e.g., via a GAMLSS-based approach [41]. This turned out to be an approach most likely suitable for in-depth analysis of a limited number of data sets. However, automatized treatment of a high number of data sets is problematic, mainly due to computational difficulties requiring manual intervention.

## Conclusion

Notos allows for robust description of CpN o/e distributions and mode detection. In the future, it seems advisable to also take other aspects into account, for example skewness and kurtosis, but also simple location measures such as the location of or distance between several modes. On the long run, DNA methylation patterns should also be investigated on sequence-level, since the reduction to a CpN o/e ratio comes along with a loss of information, such as location of the (non-)methylated regions. Such an approach would, nevertheless, require the development of suitable models, and their estimation would be by far more computationally intensive than the procedures carried out by Notos. We anticipate that already the availability of Notos will make it possible to calibrate the CpN o/e distributions with existing experimental data so that precise estimations of DNA methylation can be obtained based on Notos data.

## List of abbreviations

KDE: kernel density estimation
CpN o/e: observed to expected ratio of di-nucleotides composed of cytosine, followed by any nucleotide in 5’-3’ direction
CpG o/e: observed to expected ratio of di-nucleotides composed of cytosine, followed by guanine in 5’-3’ direction
AIC: Akaike Information criterion
BIC: Bayesian information criterion
ICL: Integrated Completed Likelihood
dbEST: database of Expressed Sequence Tags

## Ethics approval and consent to participate

No ethics approval was required for this study.

## Consent for publication

Not applicable.

## Availability of data and materials

All data generated or analyzed during this study are included in this published article and its supplementary information files. Notos is available in the Galaxy ToolShed and can also be downloaded from http://ihpe.univ-perp.fr/acces-aux-donnees/.

## Author’s contributions

IB, BA, VL, and JB performed the experiments and developed and tested the mathematical algorithms, CG designed the experiments. IB and CC implemented the software. All authors contributed to writing the manuscript. JB coordinated the work.

## Authors’ information

The authors are grateful to Rémi Emans for technical support, and David Duval and Céline Cosseau for helpful discussions. The work on this tool was initiated during a meeting that had received funding of the French-Norwegian travel program AURORA. Therefore, Notos (the son of Aurora, the goddess of dawn in Roman mythology) was chosen as name. In addition, the name refers to the simplicity (no-to-does) of the tool.

## Additional Files

### Additional file 1 — CpG o/e ratios from dbEST analyzed by Notos: data preparation output - graphics

The file ‘data cleaning species dbEST.pdf’ shows the figure produced by the data cleaning step.

### Additional file 2 — CpG o/e ratios from dbEST analyzed by Notos: data preparation output - table

The data preparation step of Notos carried out for 603 species from dbEST provides the tab-separated file ‘outliers cutoff.csv’. In the following we provide brief explanation on the content of the columns of this file. Future improvements of Notos may lead to changes, hence consult the the readme section of the galaxy interface.

- Name: name of the file analyzed
- prop.zero: proportion of observations equal to zero excluded (relative to original sample)
- prop.out.2iqr: proportion of values equal excluded if 2*·*IQR was used, relative to sample after exclusion of zeros (0 - 100)
- prop.out.3iqr: proportion of values equal excluded if 3*·*IQR was used, relative to sample after exclusion of zeros (0 - 100)
- prop.out.4iqr: proportion of values equal excluded if 4*·*IQR was used, relative to sample after exclusion of zeros (0 - 100)
- prop.out.5iqr: proportion of values equal excluded if 5*·*IQR was used, relative to sample after exclusion of zeros (0 - 100)
- used: IQR used for exclusion of outliers / extreme values
- no.obs.raw: number of observations in the original sample
- no.obs.nozero: number of observations in sample after excluding values equal to zero
- no.obs.clean: number of observations in sample after excluding outliers / extreme values

### Additional file 3 — CpG o/e ratios from dbEST analyzed by Notos: mode detection output - graphics

This file ‘modes species dbEST.pdf’ shows the graphical output from the density estimation step with activated option for the bootstrap procedure.

### Additional file 4 — CpG o/e ratios from dbEST analyzed by Notos: mode detection output - basic statistics

The density estimation step of Notos carried out for 603 species from dbEST provides the tab-separated file ‘modes basic stats.csv’. In the following we provide brief explanation on the content of the columns of this file. We are hereby using the following notation: *σ* – standard deviation, *μ* – mean, – median, *Mo* – mode, *Q*_*i*_ – the *i*-th quartile, *q*_*s*_ – the *s* % quantile. Future improvements of Notos may lead to changes, therefore consult the the readme section of the galaxy interface.

- Name: name of the file analyzed
- Number of modes: number of modes without applying any exclusion criterion
- Number of modes (5% excluded): number of modes after exclusion of those with less then 5% probability mass
- Number of modes (10% excluded): number of modes after exclusion of those with less then 10% probability mass
- Skewness: Pearson’s moment coefficient of skewness *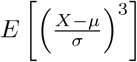*
- Mode skewness: Pearson’s first skewness coefficient 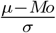
- Nonparametric skew: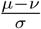
- Q50 skewness: Bowley’s measure of skewness / Yule’s coefficient 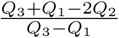
- Absolute Q50 mode skewness: (*Q*_3_ + *Q*_1_)*/*2 *- Mo*
- Absolute Q80 mode skewness: (*q*_90_ + *q*_10_)*/*2 *- Mo*
- Peak *i*, *i* = 1*,…,* 10: location of peak *i*
- Probability Mass *i*, *i* = 1*,…,* 10: probability mass assigned to peak *i*
- Warning close modes: flag indicating that modes lie too close. The default threshold is 0.2
- Number close modes: number of modes lying too close, given the threshold
- Modes (close modes excluded): number of modes after exclusion of modes that are too close
- SD: sample standard deviation *σ*
- IQR 80: 80% distance between the 90 % and 10 % quantile
- IQR 90: 90% distance between the 95 % and 5 % quantile
- Total number of sequences: total number of sequences / CpG o/e ratios used for this analysis step

### Additional file 5 — CpG o/e ratios from dbEST analyzed by Notos: mode detection output - bootstrap statistics

The optional bootstrap procedure of the density estimation step of Notos carried out for 603 species from dbEST provides the tab-separated file ‘modes bootstrap.csv’. In the following we provide brief explanation on the content of the columns of this file. Future improvements of Notos may lead to changes, thus consult the the readme section of the galaxy interface.

- Name: name of the file analyzed
- Number of modes (NM): number of modes detected for the original sample
- % of samples with same NM: proportion of bootstrap samples with the same number of modes (0 - 100)
- % of samples with more NM: proportion of bootstrap samples a higher number of modes (0 - 100)
- % of samples with less NM: proportion of bootstrap samples a lower number of modes (0 - 100)
- no. of samples with same NM: number of bootstrap samples with the same number of modes
- % BS samples excluded by prob. mass crit.: proportion of bootstrap samples excluded due to strong deviations from the probability masses determined for the original sample (0 - 100)

